# The complete genome of carbapenem-resistant *Escherichia coli* ST410 harbored *bla*_NDM-5_ isolated from reservoir water in Singapore

**DOI:** 10.1101/2021.07.28.454241

**Authors:** Yang Zhong, Siyao Guo, Glendon Ong Hong Ming, Joergen Schlundt

## Abstract

**Objective:** *Escherichia coli* ST410 with *bla*_NDM-5_ has been increasingly detected as multidrug resistance pathogens globally, even though there are very few reports of infections caused by *bla*_NDM-5_ producing *E. coli* in Singapore[1]. And significantly limit sequencing information of *bla_NDM-5_* carried *E. coli* strain from Singapore. In 2018, our group obtained a carbapenem resistance *E. coli* ST410 strain SrichA-1 isolated from reservoir water in Singapore, determined to harbor the NDM-5 gene. (BioSample Accession: SAMN18579051).

**Methods:** The susceptibility test to antimicrobials was performed with microdilution minimum inhibitory concentration (MIC) test and interpreted according to the Clinical And Laboratory Standards Institute (CLSI) -M100 standards. The genomic DNA of this strain was extracted and send for Whole-genome sequencing(WGS) with the Illumina platform. The WGS analysis was processed with the Center for Genomic Epidemiology (CGE, DTU) server.

**Results:** During the minimum inhibitory concentration (MIC) test, SrichA-1 has shown strong resistance to all the beta-lactams, including cephalosporin and carbapenem, which can not be inhibited by the clavulanic acid. Further whole genome sequencing analysis has shown that the strain harboring five beta-lactamase genes covers all class A to D, including the carbapenemase genes as *bla*_NDM-5_.

**Conclusion:** Here, we reported the complete chromosome sequence of this isolate as well as the sequence of a cycler plasmid. The pSGNDM-5 plasmid was furtherly identified to carry four beta-lactamase genes, including *bla*_NDM-5_, *bla*_CTX-M-15_, *bla*_TEM-1B_, *bla*_OXA-1_, while a *bla*_CMY-2_ was detected to be located on the chromosome.

---

*E. coli* ST410 isolate was obtained during the beta-lactamase-producing bacteria investigation from reservoir water in Singapore by Nanyang Food Technology Centre (NAFTEC). A bottle of 800 ml reservoir water was filtered with a 0.45 μm filter membrane and further enriched with the nutrition medium. The microbiota culture was then streaked on a Brilliance^™^ ESBL Agar (Thermofisher, the USA) following with purified on Thermo Scientific^™^ Mueller-Hinton (MH) Agar with 5% Sheep Blood (Thermo Scientific, the USA). The purified strain was further confirmed as carbapenem-resistant with micro-broth dilution methods using Sensititre^™^ Extended Spectrum Beta-lactamase Plate (Thermofisher, the USA). As determined by the MIC test, the isolate SrichA-1 is resistant to all the tested cephalosporin, carbapenem, and even ciprofloxacin but sensitive to gentamicin. The SrichA-1 strain has been further characterized with whole-genome sequencing (WGS) analysis. The genomic DNA was extracted with the QIAamp DNA Blood Mini Kit (Qiagen, Germany) and sequenced with the Pacbio RS II system (Pacbio, the USA). The Multilocus sequence type (MLST) was determined as ST410, serotype as H9, fimH24. Besides, 14 AMR genes were detected using CGE ResFinder 4.0 (https://cge.cbs.dtu.dk/services/ResFinder/) with the default setting[2]. The AMR genes belonged to eight classes, including macrolide, beta-lactams, phenicol, sulphonamide, quinolone, tetracycline, trimethoprim were detected. The sequence of a closed *IncF* plasmid type [F-:A1: B49] was obtained after assemble. All the resistance genes except *mdf(A)* and *bla*_CMY-2_ were found on the plasmid.

The chromosome is 4806160 bp, detected to contain 4786 genes. The closest genome found based on both cgMLST and cgSNP is strain SIEC197 (Accession: SAMN11399743) from Thailand, which carries four same beta-lactamase genes as our isolate. As the SIEC197 is a pathogen isolated from patients, a pathogenetic prediction with Virulence Finder and Pathogen Finder was also performed and identified Srich A-1 as a potential human pathogen with a probability of 0.936[3]. Virulence genes include *fyuA*, *gad*, *irp2*, and *terC* were detected on the chromosome. However, based on the phylogenetic analysis of SrichA-1, no close related clinical isolates or infection cases have been found in Singapore.

The circular plasmid, which is 84257 bp, was further aligned with other close neighbors found by NCBI Blastn from different sources and other countries. The pSGNDM-5 plasmid has shown higher identification compare to the plasmid pAMA1167-NDM-5 from Denmark, which was also reported from an ST410 isolate[4]. Except for the beta-lactamase, seven other AMR genes have been detected. Two integrases *IntI* leading gene cassette were detected on the upstream and downstream of *bla*_TEM-1_, respectively. A partial converse gene cluster contains *ISEcp1*, the *bla*_CTX-M-15_, and *WubC*, which has been reported by our group to be located on the chromosome of other *E. coli* isolates, was also find on this plasmid[5]. But the first 1375 bp of *ISEcp1* was replaced by a partial reverse transposase (Access: EIL37998.1), which may as a consequence of recombination during transfer.

To our knowledge, this is the first complete chromosome and the plasmid sequence report of the *E. coli* ST410 strain carry *bla*_NDM-5_ from Singapore.

**Fig 1.**
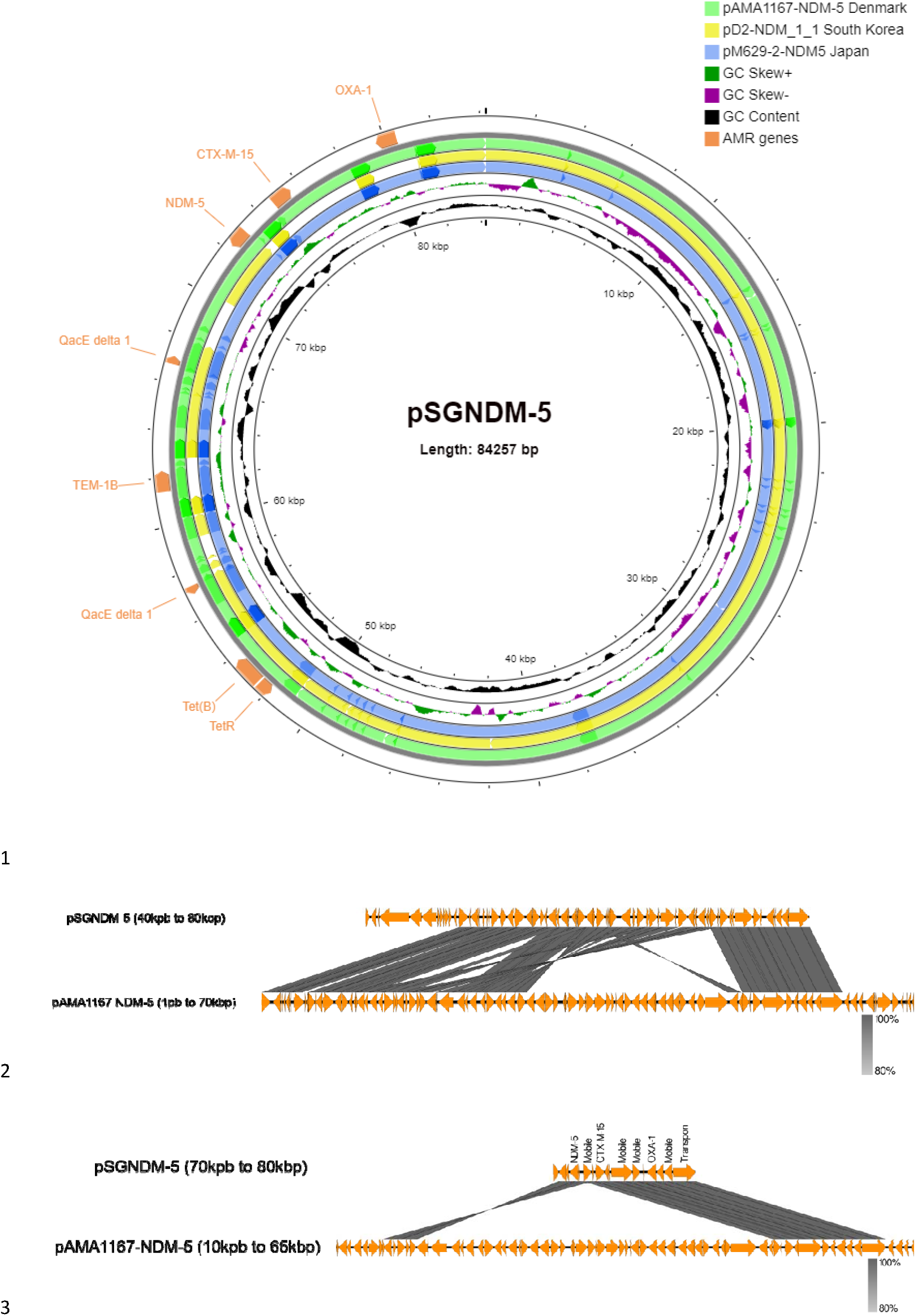
Comparison of plasmid pSGNDM-5 aligned with three similar plasmids harboring *bla*_NDM-5_. For the circulation map, the inner circle presents pSGNDM-5 as the reference sequence. Three plasmids that were also bearing *bla*_NDM-5_ were chosen for comparison and presented with different colors as middle circles. The antimicrobial genes were annotated on the outer circle as arrows. A linear alignment was performed between pSGNDM-5 and pAMA1167 NDM-5 (Denmark). Compared to pAMA1167 NDM-5 (111310 bp), pSGNDM-5 is missing a 27053 bp fragment, which presents potential recombination of the genetic environment around the *bla*_NDM-5_ gene.

**Table 1.** The minimum inhibitory concentration (MIC) tested with microdilution method.

| Antimicrobials | SrichA-1 (µg/ml) |
| --- | --- |
| Ceftriaxone | >128 |
| Meropenem | >8 |
| Cephalothin | >16 |
| Cefpodoxime | >32 |
| Ciprofloxacin | >2 |
| Cefotaxime | >64 |
| Gentamicin | <4 |
| Cefotaxime / clavulanic acid | >64/4 |
| Ampicillin | >16 |
| Ceftazidime | >128 |
| Cefazolin | >16 |
| Ceftazidime / clavulanic acid | >128/4 |
| Imipenem | 16 |
| Piperacillin / tazobactam constant 4 | >64/4 |
| Cefepime | >16 |
| Cefoxitin | >64 |

## Transparency declarations

All authors declare there is no conflict of interests.

